# Structural basis of STAT2 recognition by IRF9 reveals molecular insights into ISGF3 function

**DOI:** 10.1101/131714

**Authors:** Srinivasan Rengachari, Silvia Groiss, Juliette Devos, Elise Caron, Nathalie Grandvaux, Daniel Panne

## Abstract

Cytokine signalling is mediated by the activation of distinct sets of structurally homologous JAK and STAT signalling molecules, which control nuclear gene expression and cell fate. A significant expansion in the gene regulatory repertoire controlled by JAK/STAT signalling has arisen by the selective interaction of STATs with IRF transcription factors. Type I interferons (IFN), the major antiviral cytokines, trigger the formation of the ISGF3 complex containing STAT1, STAT2 and IRF9. ISGF3 regulates the expression of IFN–stimulated genes (ISGs). ISGF3 assembly depends on selective interaction between IRF9, through its IRF–association domain (IAD), with the coiled–coil domain (CCD) of STAT2. Here, we report the crystal structures of the IRF9–IAD alone and in a complex with STAT2–CCD. Despite similarity in the overall structure among respective paralogs, the surface features of the IRF9–IAD and STAT2– CCD have diverged to enable specific interaction between these family members, thus enabling ISGF3 formation and expression of ISGs.

## Introduction

Cytokine signalling via the JAK-STAT pathway controls the development, differentiation and regulation of cells in the immune system and is frequently dysregulated in disease ^1^. JAK-STAT signalling is mediated by four structurally related JAK kinases (JAK1, JAK2, JAK3, TYK2) and seven STAT (1, 2, 3, 4, 5a, 5b, 6) proteins^2^. A hallmark of cytokine signalling is functional redundancy and extensive pleiotropy, the ability of multiple cytokines to exert overlapping biological activities^3,4^. A critical question is how a limited number of JAK and STAT molecules enable such extensive redundancy and pleiotropy and how gene duplication and divergence among STATs contributes to specificity in cytokine signalling.

JAK–mediated tyrosine phosphorylation of STATs induces dimerization and translocation to the nucleus where STATs bind the gamma activated sequence (GAS), a palindromic 9–11 base pair (bp) DNA element, 5’–TTCN_2–4_GAA–3’ in the promoter of target genes^2^. An exception occurs in the response to type I and type III IFNs: These cytokines are rapidly induced during viral infection and stimulate activation of a complex termed ISGF3 (IFN–stimulated gene factor 3). ISGF3 contains a STAT1/STAT2 heterodimer which interacts with IRF9, a member of the IRF family of transcription factors ^5-8^. Mammals contain ten IRF paralogs that typically bind to the consensus DNA sequence 5'–AANNGAAA–3'^9-13^. As a result of STAT and IRF complex formation, ISGF3 binds to a ∼12–15 bp composite IFN–stimulated response DNA element (ISRE) 5'–G/ANGAAAN_2_GAAACT–3’. Thus the physical association of STATs with IRFs contributes to functional specificity in cytokine signalling and enables expression of IFN– stimulated genes (ISGs)^6,14^.

IRFs contain a conserved N–terminal DNA–binding domain and a C–terminal IRF–association domain (IAD; Fig. 1a). The IAD belongs to the SMAD/FHA domain superfamily^15-17^. IRF3 is the best–understood IRF family member. Signals emanating from pattern recognition receptors (PRR) activate the kinase TBK1 which phosphorylates latent IRF3^13,18-20^. This phosphorylation results in remodelling of an autoinhibitory segment in the IAD of IRF3, leading to dimerization and interaction with the transcriptional co–activators CBP/p300^21^. Although IRF9 contains a structurally related IAD, it does not share the same activation mechanism and co–activator preference with IRF3: IRF9 binds to STAT2 in both unstimulated and type I/III IFN–stimulated cells^22-24^. The interaction requires the IRF9–IAD and the coiled–coil domain (CCD) of STAT2^23^, a domain that is conserved among STAT paralogs (Fig. 1a). Thus, despite conservation of the IRF9–IAD and the STAT2–CCD among their respective paralogs, only STAT2 and IRF9 interact selectively.

**Figure 1.**
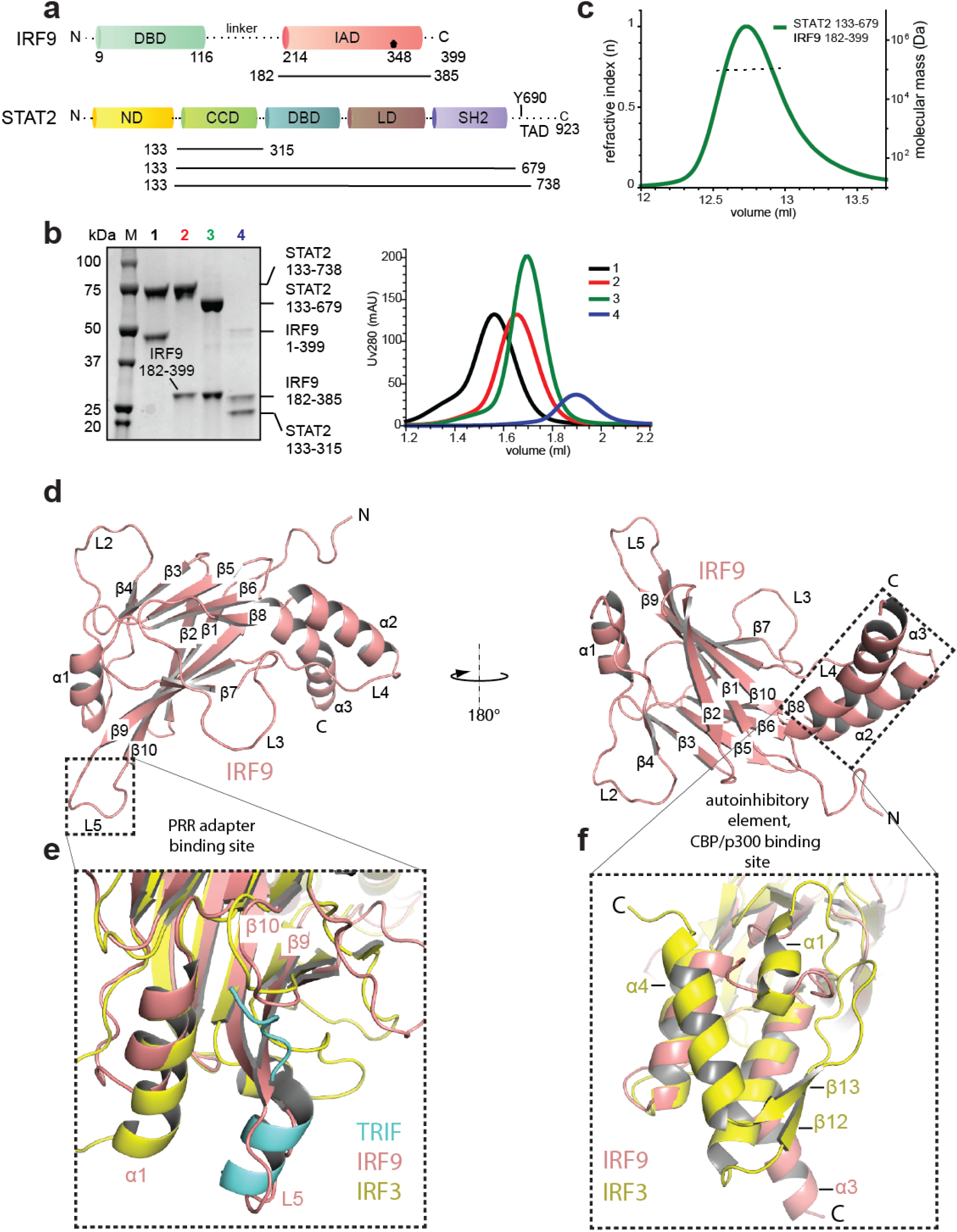
Characterization of the STAT2–IRF9 complex and structure determination of the IRF9–IAD. **a**, Domain organization of STAT2 and IRF9. DBD (DNA–binding domain), IAD (IRF–association domain), ND (N–domain), CCD (coiled–coil domain), LD (linker domain), SH2 (Src homology domain), TAD (transactivation domain). **b**, Size exclusion chromatography of different STAT2–IRF9 complexes. Fractions were analysed by SDS–PAGE. Bands corresponding to each protein are indicated. The analysed samples are 1: STAT2_133–738_–IRF9FL; 2: STAT2_133–738_–IRF9_182–399_; 3: STAT2_133–679_–IRF9_182–399_; 4: STAT2_133–315_–IRF9_182–385_. **c**, Size exclusion chromatography and multi–angle laser light scattering profile of the STAT2_133–679_– IRF9–IAD complex. The black line represents the average molecular weight across the peak. **d**, Ribbon diagram representation of the IRF9–IAD structure (red). The secondary structure elements are labelled. The model is rotated 180° between the left and right panels. **e**, Left panel: Structural overlay of IRF9 (red) and IRF3 (yellow). The extended strands β9 and β10 of IRF9 block access of the PRR adaptor TRIF (blue). **f**, Structural overlay of IRF9 and autoinhibited IRF3–IAD (yellow). The autoinhibitory element of IRF3 absent in IRF9

To explore the molecular basis that enables selective IFR9–STAT2 interaction, we have determined the crystal structure of the IRF9–IAD in isolation and in complex with STAT2. As expected, the IRF9–IAD is closely related to IRF3. However, IRF9 lacks the structural elements involved in IRF3 autoinhibition thus explaining the different activation, co–factor and oligomerisation requirements. IRF9 binds through a hydrophobic patch on the convex side of the IAD to the tip of the STAT2–CCD. The binding interface is conserved in IRF9 and STAT2 orthologs but is divergent in other IRF and STAT paralogs. Point–mutations in the conserved interface disrupt STAT2–IRF9 interaction *in vitro* and abolish ISGF3 activity in cells. Overall, our data reveal the molecular basis for selective IRF9 and STAT2 interaction. Comparative structural and biochemical analyses yield insights into how gene duplication, evolutionary drift and natural selection of STATs and IRFs resulted in expansion of the gene regulatory repertoire in cytokine signalling.

## Results

### Biochemical basis for STAT2–IRF9 interaction

STAT2 contains an N–domain (ND), a CCD, a DNA–binding domain (DBD), a linker domain (LD), an SH2 domain and a C–terminal transactivation domain (TAD) (Fig. 1a). Phosphorylation on Y690 enables dimerisation of STAT2^5,25^. IRF9 contains a N–terminal DBD and a C–terminal IAD connected by a flexible linker (Fig. 1a). Previous data show that the region spanning amino acids 148–324 of STAT2 and amino acids 217–377 of IRF9 are required for complex formation^8,23,26^. To further delimit the interacting regions, we co–expressed STAT2_133–738_ and full–length IRF9 (IRF9FL) and purified the complex to homogeneity. We found that IRF9FL co– purified with STAT2_133–738_ (Fig. 1b, lane 1) and that a C–terminal fragment of IRF9 spanning 182–399 was sufficient for STAT2 binding (Fig. 1b, lane 2). Limited proteolysis using Trypsin digestion resulted in C–terminal truncation of STAT2 at position 679 as revealed by LC–MS (Extended Data Fig.1a). This shows that the region beyond the SH2 domain is flexibly attached and is not required for complex formation. Co–expression of STAT2_133–679_ with IRF9_182–399_ resulted in a stable complex (Fig. 1b, lane 3). Analysis by size exclusion chromatography coupled to multi–angle laser light scattering (SEC–MALLS) showed a monodisperse complex with a molecular mass of 86.8 ± 0.7 kDa, in agreement with the expected mass of a 1:1 heterodimer (Fig. 1c). Chymotryptic digestion resulted in cleavage of the 14 terminal residues of IRF9, further delimiting the interacting region (Extended Data Fig.1b). Co–expression of STAT2_133–679_ or of the coiled–coil segment STAT2_133–315_ with IRF9_182–385_ resulted in stable and monodisperse heterodimeric complexes (Fig. 1b, lane 4). We conclude that the minimal regions required for complex formation comprise STAT2_133–315_ and IRF9_182–385_. Considering that ISGF3 contains a single copy of STAT1 and STAT2^27^, we propose an overall 1:1:1 stoichiometry.

### The IRF9–IAD lacks the autoinhibitory element and the PRR adapter binding site

We obtained crystals of the IRF9_182–385_ IAD, which diffracted to 1.9Å resolution, and determined the structure by molecular replacement. Our final model includes residues 197–385 of IRF9 (Table 1). Like other IAD domains^15,17,28^, IRF9–IAD has a MH2 domain fold - a central β– sandwich core (β1–β10) flanked by a set of helices and loops (Fig. 1d). The domain has a crescent–like shape with a two–helix bundle (α2, α3) on one end, where the N– and C– termini are located, and helix α1 on the other end. While the β–sandwich core of the IAD domain is conserved among IRFs, the connecting secondary structures and loop regions vary: For example, IRF9 lacks the C-terminal tail of IRF3, constituted by the strands β12, β13 and helices α1 and α4, which autoinhibit IRF3 in the latent form (Fig. 1e). This C–terminal element of IRF3 undergoes a conformational change from a buried, autoinhibitory configuration to an extended coil structure that leads to formation of a domain– swapped dimer upon TBK1–mediated phosphorylation (Extended Data Fig. 1c)^21,29^. The rearranged IAD of IRF3 thereby exposes a hydrophobic binding site for the coactivators CBP/p300 (Extended Data Fig. 1c). The missing autoinhibitory element of IRF9 renders the hydrophobic residues of helices α2 and α3 surface exposed (Fig. 1d-e).

**Table 1:** Xray data collection, phasing and refinement statistics.

| Parameters | IRF9-IAD | STAT2-IRF9 |
| --- | --- | --- |
| **Data collection** |  |  |
| Space group | $P3_221$ | $P2_1$ |
| Cell dimensions |  |  |
| $a, b, c$ (Å) | 76.77, 76.77,<br>85.6 | 30.59, 123.96,<br>50.99 ($\beta=92.1^\circ$) |
| Wavelength (Å) | 0.966 | 0.972 |
| Resolution (Å) | 42.8–1.9 | 27.0–2.7 |
| Total reflections | 66664 | 31975 |
| Unique reflections | 22819 | 9943 |
| $R_{\text{sym}}$ or $R_{\text{merge}}$ | 0.03 (0.94)* | 0.33 (2.35)* |
| $I / \sigma I$ | 14.5 (1.0) * | 4.2 (1.0)* |
| Completeness (%) | 97.2 (82.2) * | 95.5 (75.8)* |
| Redundancy | 2.9 (2.1) * | 3.2 (3.1) |
| Wilson $B$-factor (Å<sup>2</sup>) | 42.7 | 52.34 |
| **Refinement** |  |  |
| $R_{\text{work}} / R_{\text{free}}$ | 20.6 / 24.2 | 22.9 / 29.6 |
| No. of atoms | 2970 | 2769 |
| Protein | 2933 | 2698 |
| Ligand | 5 | N/A |
| Solvent | 32 | 71 |
| $B$-factors (Å<sup>2</sup>) | | |
| Protein | 70.3 | 54.6 |
| Ligand | 68.3 | N/A |
| Solvent | 55.7 | 28.8 |
| R.m.s deviations |  |  |
| Bond lengths (Å) | 0.014 | 0.009 |
| Bond angles (°) | 1.495 | 0.98 |
| Ramachandran distribution |  |  |
| Favoured | 93.05 | 94.82 |
| Allowed | 6.95 | 5.18 |
| Outliers | 0.0 | 0.0 |

To calculate $R_{\text{free}}$ , 5% of the reflections were excluded from the refinement. $R_{\text{sym}}$ is defined as $R_{\text{sym}} = \sum_{\text{hkl}} \sum_i |I_i(\text{hkl}) - \langle I(\text{hkl}) \rangle| / \sum_{\text{hkl}} \sum_i I_i(\text{hkl})$ . \*Values in parentheses are for the highest-resolution shell. N/A; not-applicable

In IRF9, the region comprising the helix α1, strands β9 - β10 and the connecting loop L5 correspond to the binding of the pLxIS motif of phosphorylated PRR adaptors such as STING, MAVS, TRIF or alternatively, the phosphorylated C-terminal tail to form a loop-swapped dimer in IRF3 (Fig. 1e, Extended Data 1e)^21^. The extended structure of the strands β9, β10 and the connecting loop L5 in IRF9, abolish access to the PRR adaptor binding/dimerization site (Fig. 1e). The lack of the PRR adaptor binding site and of the autoinhibitory/dimerisation element thus explains the different activation and oligomerisation properties of IRF9 as compared to IRF3 and other IRF family members.

### STAT2 binds to the convex surface on the IRF9-IAD

To identify regions of IRF9 that are potentially involved in STAT2 binding, we compiled sequence alignments of IRF9 from divergent vertebrates and mapped the amino acid conservation onto the IAD structure (Fig. 2a). We focused on conserved residues of the convex surface of the β-sandwich (IF1) and helix α2 (IF2) and on residues of helix α2 and α3 (IF3), that are involved in CBP/p300 binding in IRF3. We mutated these three putative interfaces in IRF9 and tested binding to STAT2 using a Ni^2+^-affinity pull-down assay (Fig. 2b). Mutations in IF1 reduced or completely abolished IRF9 interaction with STAT2 (Fig. 2b). These include IF1-A (R236E, F283A), IF1-B (R236E, L274A, F283A) or IF1-C (L233A, R236E, L274A F283A). In contrast, mutations in IF2 (R324E, D325K, Q331E, Q333E, P335S) or IF3 (L326A, F330A, I376A) retained STAT2 binding activity (Fig. 2b). Thus, our data suggest that STAT2 binds to the convex surface of the IRF9-IAD through residues in IF1.

**Figure 2.**
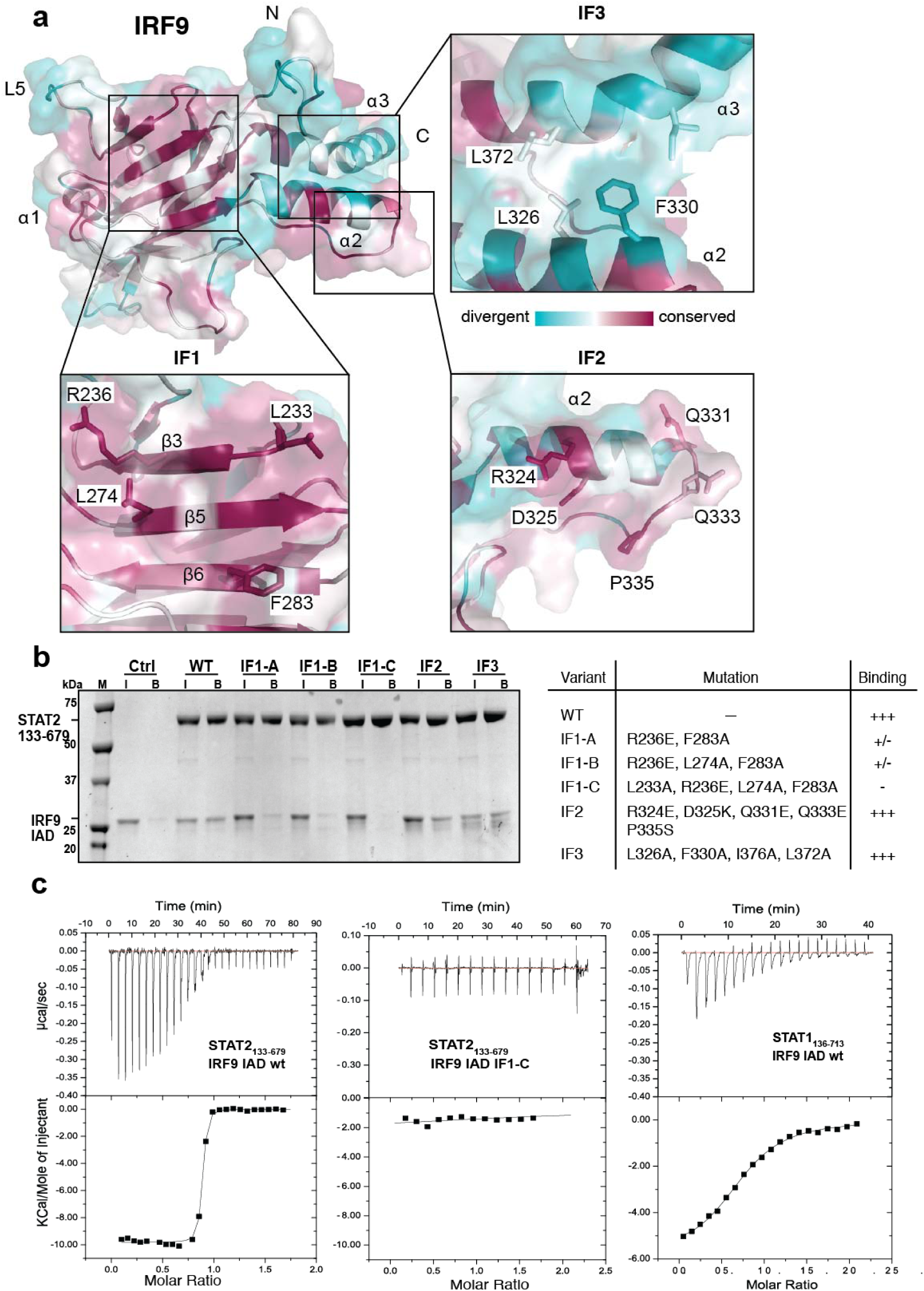
Characterization of the STAT2 binding interface on IRF9. **a**, The IRF9–IAD is coloured according to amino acid conservation among vertebrate IRF9 orthologs. Magenta = well conserved, light blue = highly variable; for sequence alignment, see Extended Data Fig. 2. Panels: zoom–in onto the conserved surfaces boxed in **(a)**, showing the underlying amino acid residues that were analysed by mutagenesis. **b**, Purified His_6_–tagged STAT2_133–679_ was incubated with IRF9–IAD variants containing point mutations in the different conserved surfaces. The resulting complexes were incubated with Ni^2+^–affinity resin, and bound proteins analysed by SDS–PAGE. Table: Summary of mutants analysed: Mutants in the IF1 interface reduced or abolished STAT2 binding. Mutation of residues in the IF2 or IF3 interface did not interfere with STAT2 binding. **c**, Isothermal titration calorimetry (ITC) binding curves for interaction between STAT2_133–679_ and IRF9–IAD, STAT2_133–679_ and IRF9–IAD IF1–C or STAT1_136–713_ and IRF9– IAD.

To analyse the impact of these mutants on STAT2–IRF9 interaction more quantitatively, we measured the equilibrium binding isotherms by isothermal titration calorimetry (ITC). STAT2_133–679_ bound to the IRF9–IAD exothermically, with an equilibrium dissociation constant (K_d_) of 10 nM (Fig. 2c). As expected, the IF2 and IF3 mutants retained close to wild–type binding affinity (Table 2). The double mutant IF1–A and the triple mutant IF1–B had barely detectable STAT2 binding activity while binding of the quadruple mutant IF1–C was completely abolished (Fig. 2c, Extended Data Fig. 3a,b). Thus, our data indicates that the conserved residues in IF1 synergistically contribute to STAT2 binding. Previous studies have implicated a potential interaction between STAT1 and the C–terminus of IRF9 but the relevance for ISGF3 signalling has remained unclear^8,23,30^. We analysed the interaction between STAT1_136–713_ and the IRF9– IAD by ITC and found an equilibrium dissociation constant, K_d_, of 5 μM (Fig. 2c, Table 2). The interaction with STAT1 is also mediated by the IF1 interface of IRF9 as the IF1–C mutant completely abolished STAT1 binding (Extended Data Fig. 3c). The 500–fold higher binding affinity for STAT2 likely explains why IRF9 constitutively interacts with STAT2 but not STAT1 in ISGF3 signalling^23^.

**Table 2:** Summary of ITC data.

| **STAT variant** | **IRF9 variant** | **Mutation(s)** | **K<sub>d</sub> (nM) \*** |
| --- | --- | --- | --- |
| STAT2 WT | IRF9 WT | – | 10 nM |
|  | IF1 -A | R236E, F283A | n.b. |
| STAT2 WT | IF1 -B | R236E, F283A, L274A | n.b. |
|  | IF1 -C | R236E, F283A, L274A, L233A | n.b. |
| STAT2 WT | IF2 | R324E, D325K, Q331E, Q333E, P335S | 11 nM |
| STAT2 WT | IF3 | L326A, F330A, I376A | 17 nM |
| STAT2 F174D | IRF9 WT | F174D | n.b. |
| STAT2 V171E | IRF9 WT | V171E | 2.5 μM |
| STAT1 WT | IRF9 WT | – | 5 μM |
| STAT1 WT | IF1 -C | R236E, F283A, L274A, L233A | n.b. |
| STAT1 E169V | IRF9 WT | E169V | 116 nM |

\* n.b.: no binding

**Figure 3.**
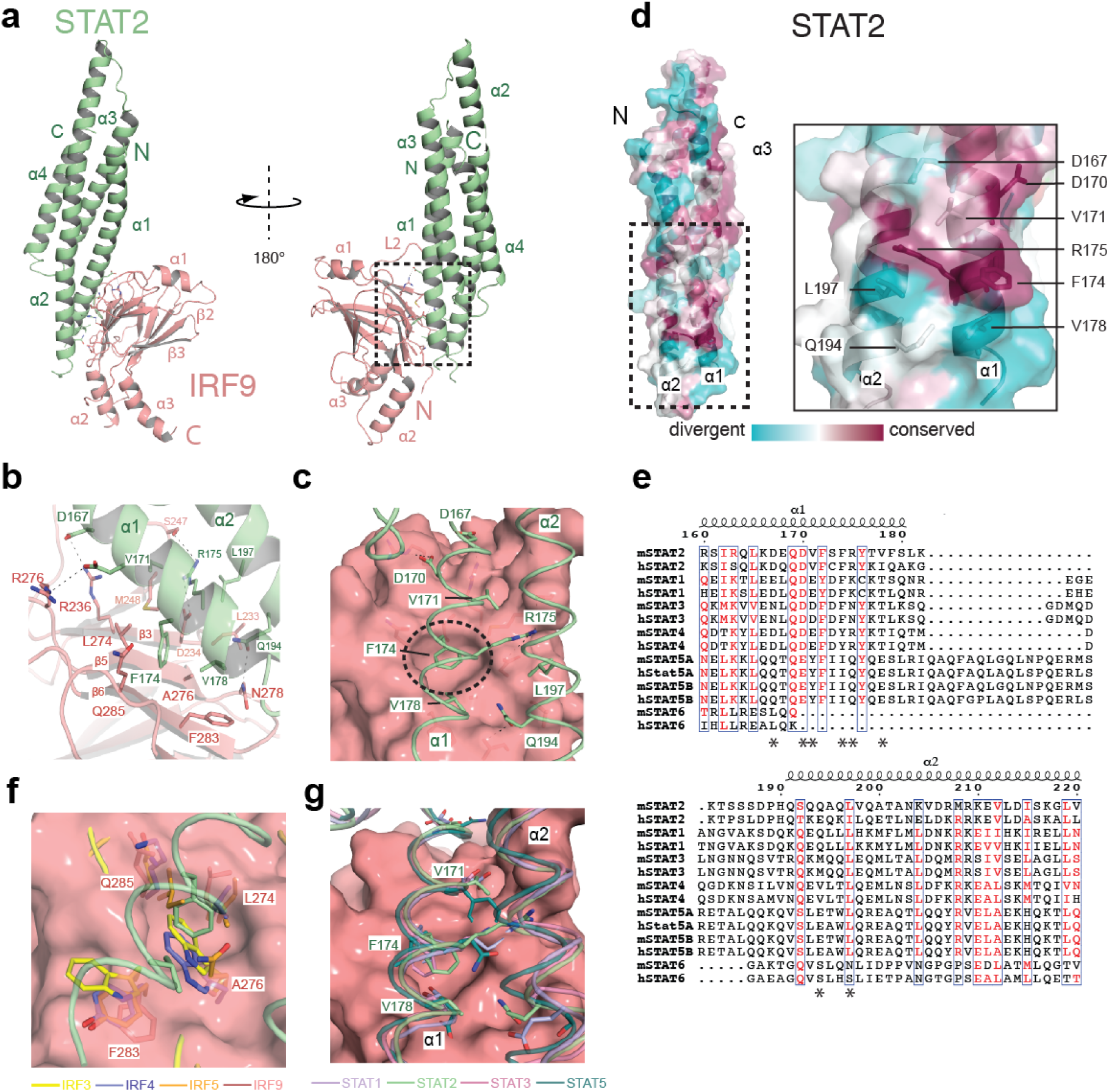
Structure of the STAT2–IRF9 complex. **a**, STAT2 (green) and IRF9 (red) are shown as ribbons. The model is rotated 180° between the left and right panels. **b**, Expanded view of the STAT2:IRF9 interface. Residues in the interface are shown with nitrogen in blue and oxygen in red. Carbon atoms are coloured according to residue location. **c**, Surface drawing of IRF9 with the recognition helices α1 and α2 of STAT2 shown as coils. Residues of STAT2 in the interface are indicated and the key anchoring residue F174 is circled. **d**, The STAT2–CCD is coloured according to amino acid conservation among vertebrate STAT2 orthologs. Magenta = well conserved, light blue = highly variable. Panel: zoom–in onto the conserved surface patch showing the underlying amino acid residues. **e**, Sequence alignment of mouse and human STAT family members. Residues of STAT2 involved in IRF9 binding are indicated (*). **f**, Surface representation of IRF9 superposed with IRF3, 4 and 5. A276 of IRF9 which is critical for accommodation of F174 of STAT2 is replaced by bulky side chains in in IRF3, 4 and 5. STAT2 is represented as coils. **g**, Structural overlay of STAT1, STAT3 and STAT5 onto STAT2. The secondary anchoring residues V171 and V178 are replaced by charged or bulky residues in other STATs.

### Structure of the STAT2–IRF9 complex

We obtained crystals of IRF9–IAD_182–385_ in complex with the STAT2–CCD_133–315_, which diffracted to 2.7 Å resolution, and we determined the structure by molecular replacement (Table 1). The production of diffraction–quality crystals required surface entropy reduction by introducing mutations Q242A and K243A in an exposed surface loop of STAT2. We located a single copy of the STAT2–CCD in the asymmetric unit bound to the convex side of the IRF9– IAD (Fig. 3a). The STAT2–CCD contains a four helix bundle with a left–handed twist similar to that of other STAT proteins including STAT1, 3 and 5 with which it superposes with a root– mean–square–deviation (rmsd) of between 2.7–2.9 Å for 151 aligned Cα atoms (Fig. 3g). As in other STATs, with the exception of STAT6, helices α1 and α2 extend beyond the core of the four helix bundle of the CCD. The β–sandwich core of the IRF9–IAD binds to this extension of α1 and α2 of STAT2 and buries 1040 Å^2^ of surface area (Fig. 3a). The most significant contribution to the binding interface is made by contacts between α1 of STAT2 which packs against a shallow binding groove on the surface of the IRF9–IAD. STAT2 is the most divergent STAT family member, apparently due to frequent viral targeting which has driven STAT2 divergence^31^. Inspection of amino acid conservation among 21 vertebrate STAT2 orthologs revealed a conserved patch on the surface of α1 (Fig. 3b). This patch, comprising residues D167, D170, V171, F174 and R175, faces directly into the STAT2–IRF9 binding interface. The lead anchoring residue is F174 which buries the largest solvent–accessible surface area (142 Å^2^) upon complex formation. F174 binds into a pocket on the surface of IRF9 (Fig. 3c).

Unsurprisingly, a F174D mutation completely abolished complex formation with IRF9 (Extended Data Fig.3d). V171 and V178 are secondary anchoring residues and project into two hydrophobic pockets on IRF9. Additional interacting residues of STAT2 include D167, D170 and R175 from helix α1 and Q194, and L197 from helix α2 which form hydrophobic interactions, van der Waals contacts and salt bridges with conserved surface residues in IRF9 (Fig. 3c).

On IRF9, the interacting amino acids are L233, R236, S247, M248, L274, A276, N278, F283 and Q285. The four conserved IRF9 amino acids L233, R236, L274 and F283, that are required for STAT2 binding (Fig. 2b,c), contribute a large fraction of the overall buried surface area in this interface. Residues L274, F283 together with A276, D234 and M278 line the pocket that binds F174 of STAT2.

The IRF9–IAD is closely related to those of IRF3–8 and superposes with the IADs of IRF3– IRF5 with an rmsd of between 2.7–2.9Å for 170 aligned Cα atoms (Extended Data Fig.3g). Key residues that are involved in STAT2 interaction are conserved in IRF9 orthologs and explain binding selectivity. A critical amino acid residue of IRF9 is A276. A residue with a short side chain at this position is required for formation of the binding pocket for F174 of STAT2. The methyl group of A276 makes hydrophobic contacts with the side chain of F174 of STAT2 and is completely buried upon complex formation. A276 is replaced by a bulky amino acid in other IRFs (Fig. 3f, Extended Data Fig.2). As a result, the binding pocket is absent in other IRFs thus preventing association of STAT2 (Extended Data Fig.3g). Hydrophobic pockets of IRF9 that accommodate the secondary anchoring residues V171 and V178 of STAT2 are also missing in other IRFs (Extended Data Fig.3g). Comparison of IRF9 in the free and STAT2–bound state shows the conformational rearrangements of IRF9 upon STAT2 binding (Extended Data Fig.3h). Loop L2 is displaced to accommodate STAT2 on the shallow binding groove of IRF9. This binding groove is occluded in other IRFs due to a longer L2 loop that projects into the binding site (Extended Data Fig.2, Extended Data Fig.3h). In short, despite sharing a similar overall structure with other IRFs, the surface features of the IRF9–IAD have diverged to enable specific recognition of STAT2.

Sequence comparison of STATs shows that F174, the lead–anchoring residue for IRF9 binding of STAT2, is conserved in STAT1 and STAT3 but divergent in other paralogs (Fig. 3e). This likely explains why STAT1 can interact with IRF9^8^. The 500–fold lower binding affinity of STAT1 for IRF9 (Fig. 2c) is due to divergence of the secondary anchoring residues V171 and V178. In particular, the negatively charged amino acid residues that replace V171 in STAT1 and STAT3 would clash with hydrophobic residues of IRF9. To demonstrate the importance of this interaction, we introduced a V171E mutation in STAT2 and found a 250–fold increased K_d_ (from 10 nM to 2.5 μM) for IRF9 binding (Extended Data Fig. 3e). Conversely, introduction of E169V into STAT1 resulted in a more than 40–fold reduced K_d_ (from 5 μM to 116 nM; Extended Data Fig. 3a). Together, while association between STAT2 and IRF9 is driven by F174 of STAT2, the secondary anchor residues and in particular V171 are critical for high binding affinity and specificity of the interaction.

### Effects of point mutations on ISGF3 activity

To further analyse the physiological relevance of the IRF9–STAT2 interface for ISGF3 function in cells, we introduced mutations into FLAG–tagged IRF9 and tested their interaction with HA-STAT2 in HEK293 cells and the impact on ISGF3 activity using an *IFIT1*promoter luciferase reporter (*IFIT1*prom–Luc) in IRF9^−/−^ /IRF−3^−/−^ MEFs. While IRF9wt was able to robustly Co-immunoprecipitate with STAT2 (Fig. 4a, lane 2), the interaction was reduced to background levels with the IRF9 IF1–A, IF1–B or the IF1–C mutants (Fig. 4a, lanes 3-5). IRF9 IF1-D, a variant containing the mutations L233A, L274A, F283A also did not bind STAT2 (Fig. 4a, lane 6). As the IF1 mutations were sufficient to disrupt the interaction between full-length STAT2 and IRF9 *in vivo*, we conclude that the identified binding interface is necessary and sufficient for this interaction.

**Figure 4.**
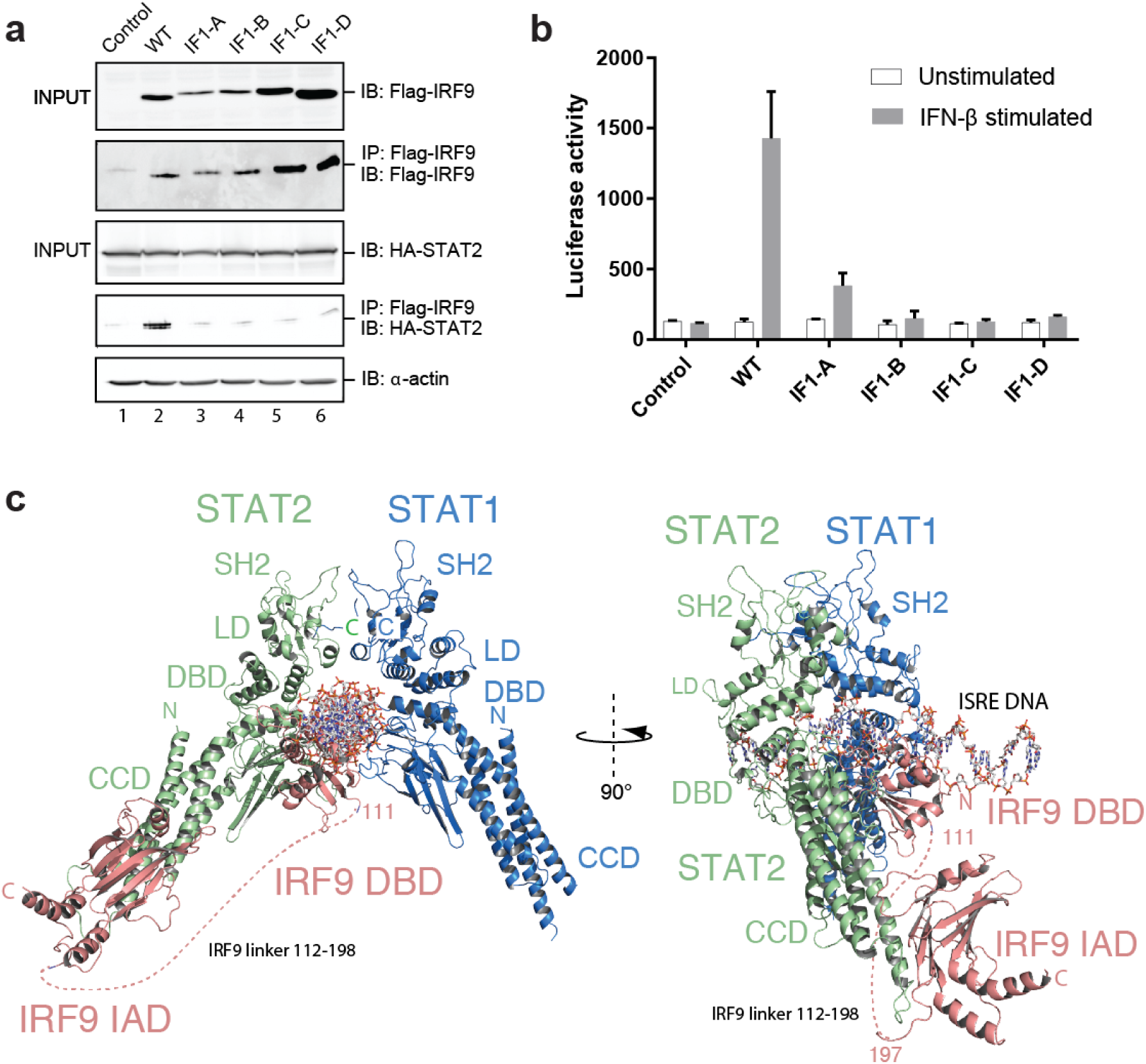
Effects of point mutations in the STAT2–IRF9 interface on ISGF3 activity. **a**, Co-immunoprecipitation of IRF9 variants with STAT2: HEK293 cells were co-transfected with FLAG-IRF9 WT or IF1 mutants and HA-STAT2. The empty FLAG vector was used as a control. Immunoprecipiation (IP) was performed with anti-FLAG beads and immunoblotted (IB) for FLAG_IRF9 or HA-STAT2. Actin concentration was used as a loading control for INPUT conditions. Two independent experiments were performed with consistency and one representative example is shown. **b**, MEF IRF9^−/−^/IRF−3^−/−^ cells were co–transfected with the *IFIT1*prom– *Firefly* luciferase reporter and Renilla luciferase plasmids along with the indicated Flag–IRF9 expression plasmids. Cells were treated with 200U/mL IFNβ prior to assaying for luciferase activities expressed as *Firefly/Renilla* ratio. Three independent experiments were performed and the mean value ± SEM are shown. **c**, Model of the ISGF3 complex bound to a ISRE DNA sequence 5’–GGGAAATGGAAACT-3. A STAT1 dimer bound to a GAS DNA sequence (1BG5) was positioned on the first 5’–GGAA–3’ repeat. A homology model of the IRF9 DNA–binding domain (DBD) was positioned on the second GAAA repeat based on the structure of DNA–bound IRF3 (1T2K). The STAT2–IRF9 complex was overlaid with one copy of STAT1 of the STAT1 dimer to obtain the final model. A dashed line indicates the flexible amino acid linker spanning residues 112–198 between the IAD and DBD of IRF9. CCD, coiled– coil domain; LD, ligand binding domain; IAD, IRF association domain.

The luciferase reporter assay showed, as expected, IFNβ induced *IFIT1*prom–Luc activity in cells reconstituted with FLAG–IRF9wt (Fig. 4b, Extended Data, Fig. 4a). However, cells expressing the IF1 mutants exhibited only partial (IF1–A and IF1–B) or no (IF1–C) *IFIT1*prom– Luc activity in response to IFNβ. The progressive loss of activity observed with IRF9 mutants, IF1–A> IFI1–B>IFI1–C correlates with our *in vitro* binding studies (Fig. 2b). Not surprisingly, the IF1–D variant, also did not show *IFIT1*prom–Luc activity (Fig. 4b). Overall, our results show that mutations to the conserved surface patch disrupt IRF9–STAT2 interaction and abolish ISGF3 function in cells.

### Solution structure of STAT2-IRF9 and model of the ISGF3 complex

We also examined the architecture of the STAT2–IRF9 complex in solution by cross– linking/mass spectrometry (XL–MS) and size–exclusion chromatography in line with small– angle X–ray scattering (SEC–SAXS). We focused on a STAT2–IRF9 complex containing an extended STAT2_133–679_ construct spanning from the CCD to the SH2 domain. XL–MS showed intermolecular crosslinks between K183 in STAT2–CCD and K318 or K381 of IRF9–IAD, in agreement with our structure (Extended Data Fig. 4b). The SAXS scattering curve allowed us to determine the radius of gyration (R_g_) of 4.37±0.02 nm while the distance distribution function *p(r)* showed a curve characteristic of an elongated particle with a maximum diameter of 14.4±0.3 nm (Extended Data Fig. 4c,d and inset). These data allowed rigid body modelling of STAT2_133–_ 679 spanning from CCD to SH2 domain in complex with the IRF9–IAD (Extended Data Fig. 4b). The resulting model represented a good fit to the experimental data (Extended Data Fig. 4c). Based on this model, together with available structures of DNA–bound STAT1 and IRF dimers, we propose a composite model for ISGF3, containing a 1:1:1 complex of STAT1:STAT2 and IRF9 bound to ISRE DNA (Fig. 4c). In this model, the IRF9–IAD is attached to the STAT2– CCD and is linked by a flexible linker spanning residues 112–198 to the IRF9–DBD. STAT1 occupies the first and the IRF9–DBD the second 5’–GAAA–3’ repeat in the major groove on opposite sides of the DNA. This positions STAT2 upstream on the same face of the ISRE DNA as the IRF9–DBD, thus explaining DNA–binding specificity of ISGF3.

## Discussion

STATs and IRFs have arisen by gene duplication, evolutionary drift and natural selection under constant selective pressure from pathogen infection^13,32,33^. The direct interaction between STAT2 and IRF9 is critical for the function of ISGF3 and the antiviral response^23^. Our studies show that the overall architecture of these domains is similar to that of other STAT and IRF paralogs. However there are several important adaptations: The IRF9–IAD is missing the regulatory apparatus that is used for IRF autoinhibition in the latent form and that in the activated state enables IRF dimerization and interaction with the transcriptional co–activators CBP/p300 ^15,17^. In addition, the PRR adaptor–binding site that enables PRR–signal–dependent activation of IRF3 is absent in IRF9. IRF9 binds to the tip of the STAT2–CCD using the convex surface of the β– sandwich core of the IAD domain. While the same surface is available in other IRF paralogs, amino acid substitutions at the key anchoring points account for the preferential STAT2 binding. Together, these adaptations explain why IRF9 binds constitutively and selectively to STAT2^23^.

The STAT2–CCD is closely related to that of that of other STATs, with the exception of STAT6, which is missing the IRF9 binding site entirely. The lead–anchoring residue of STAT2, F174, is conserved in STAT1 and STAT3. Amino acid residues at this position are critical for stabilizing an ‘antiparallel’ and apparently inactive dimer structure of STAT1 (F172) and STAT5 (I174) and thus play an important role in the STAT activation and inactivation cycle^34,35^ (Fig. 3e). The secondary anchoring residues V171 and V178 of STAT2 are divergent. Due to this divergence, STAT1 binds with 500–fold weaker affinity to IRF9 as compared to STAT2, contributing to differential binding energetics. Previous studies have reported an interaction between STAT1 and IRF9 under certain experimental conditions^8^, which is likely due to this residual binding affinity between STAT1 and IRF9. However other experiments could not reproduce these results^23^. Nevertheless, STAT1 and IRF9 appear to interact functionally on certain promoters even in the absence of STAT2 ^36,37^. Overall, our data shows that IRF9 selectively and constitutively interacts with STAT2, but only weakly with STAT1.

Together, functional divergence of the STAT and IRF paralogs is due to amino acid substitutions at key anchoring points in the binding interface that enable select family members (STAT2– IRF9) to interact with high affinity and thereby restricting interaction between other STAT and IRF paralogs. Our study thus provides evidence for how gene duplication and divergence can contribute to the evolution of tightly integrated systems by slight structural variation of key amino acids of primordial variants. These variations resulted in new protein–protein interactions that enable a significant increase in the regulatory repertoire of the mammalian cytokine response.

## Methods

### Constructs, Expression and Purification

Sequences encoding murine IRF9, STAT1 and STAT2 and any variants described in the text, were amplified by PCR from cDNA clones, and inserted between the *NcoI* and *HindIII* sites in the first open reading frame (ORF) of pETDuet vector. Co–expression was achieved by cloning respective cDNA into the second ORF between *NdeI* and *KpnI* sites. Mutants of IRF9 were generated using the QuikChange site–directed mutagenesis kit (Agilent). Sequences of all expression constructs were confirmed by DNA sequencing. Individual proteins and protein complexes were expressed in *E.coli* BL21 (DE3) cells and induced with IPTG. Upon reaching an OD_600nm_=0.6, *E.coli* cultures were shifted from a temperature of 37 °C to 16 °C, for 12 hours. Cells were pelleted at 9000 *g* (JLA–8.1, Beckman) followed by resuspension in buffer A (20 mM HEPES pH 7.5, 300 mM NaCl and 0.5 mM TCEP) containing 10 mM imidazole supplemented with protease inhibitors (Roche), and lysed using a microfluidiser (Microfluidics). Lysates were centrifuged at 24000 *g* for 30 minutes, using a JA–25.5 rotor (Beckman), and the resultant supernatant passed over Ni^2+^–conjugated IMAC sepharose resin (GE Healthcare). Columns were subsequently washed using buffer A containing 20 mM imidazole, and eluted with buffer A containing 500 mM Imidazole. For all purifications involving His–TEV proteins the Ni^2+^ eluate was incubated overnight at 4 °C with His–tagged TEV protease, during dialysis against buffer A with 20 mM imidazole. Cleaved tags, TEV protease and uncleaved protein were removed by subtractive purification over Ni^2+^ resin. All proteins were further purified by SEC using a HiLoad 16/60 Superdex 200 prep–grade column (GE Healthcare) equilibrated in buffer A.

### Limited proteolysis

To identify stable variants of STAT2 and IRF9 we performed limited proteolysis by Trypsin and α–chymotrypsin. The purified STAT2_133–738_– IRF9_182–399_ complex (120 μg) was titrated against decreasing concentrations of Trypsin ranging from a ratio of 1:125 to 1:1000 (w/w) and incubated for up to 30 minutes at 4 °C. The samples were then analysed by SDS–PAGE, acid hydrolysis and mass spectrometry. The stable STAT2_133–679_– IRF9_182–399_ (120 μg) complex identified by trypsin digestion was further proteolysed by α–chymotrypsin for 72 hours at a ratio of 1:5000 (w/w) at 4 °C. The fragments were analysed as above.

### Crystallisation

Initial trails of IRF9_182–385_ yielded crystals with inherent pathologies like twinning, translational pseudo-symmetry and high copy number. To overcome these problems, we mutated surface residues to induce alternate crystal packing. An E348A mutant yielded crystals in 0.1 M HEPES pH 6.8, 1.5 M Ammonium phosphate monobasic and 0.1 M Ammonium sulphate that diffracted to a minimum Bragg spacing of 1.9 Å. Production of diffraction quality crystals of the STAT2- IRF9 complex also required mutation of surface residues Q242A and K243A of STAT2 and of E347A and E348A of IRF9. Crystals of the STAT2-IRF9 complex (10 mg ml^−1^) were obtained in 0.2M Potassium formate and 20% PEG3350. These were further optimised by microseeding, yielding crystals that diffracted to 2.7 Å resolution. 25% glycerol was used as a cryo-protectant before flash cooling in liquid nitrogen. IRF9-IAD diffraction data were collected at 100 K at an X-ray wavelength of 0.966 Å at beamline ID30A-1 (MASSIF-1) of the European Synchrotron Radiation Facility (ESRF, Grenoble, France) with a Pilatus 6M-F detector^38^. Diffraction data for crystals of the STAT2-IRF9 complex were collected at a wavelength of 0.972 Å at the beamline ID23-1 of ESRF with a Pilatus 6M-F detector. Indexing and scaling of the data was performed with XDS and XSCALE^39^. The structure was solved by molecular replacement using the IRF5- IAD (PDB; 3DSH) structure as a search model^40^. STAT2-IRF9 data was indexed with XDS and scaled using AIMLESS^41^. The structure of the IRF9-IAD and STAT1-CCD (PDB; 1BF5) were used as search models for molecular replacement. Final models were produced by iterative rounds of automatic and manual model building and refinement, using Coot and PHENIX^42,43^. The final IRF9-IAD model contained residues 197–385 and was refined to a resolution of 1.9 Å with an R_work_ and an R_free_ of 20.6% and 24.2%, respectively (Table 1). Analysis of the refined structure in MolProbity showed that there are no residues in disallowed regions of the Ramachandran plot. The MolProbity all atom clash score was 4.7 placing the structure in the 97th (best) percentile of structures (n = 773) refined at comparable resolution^44^. The final STAT2–IRF9 model contained residues 141–315 of STAT2 and 206–376 of IRF9 and was refined to a resolution of 2.7 Å with R_work_ and an R_free_ of 22.9% and 29.6%, respectively (Table1). The MolProbity all atom clash score was 0.37 placing the structure in the 100^th^ (best) percentile of structures (n = 175). Figures displaying the structures were generated using PyMol^45^.

### SEC–MALLS and SAXS analysis

SEC was performed at 20 °C with a Superdex 200 10/300 GL column (GE Healthcare) equilibrated in buffer A. Fifty μl of STAT2_133–679_/IRF9–IAD was injected at 7 mg/ml and the sample eluted at a flow rate of 0.5 ml/min. Multi angle laser light scattering was recorded with a laser emitting at 690 nm using a DAWN–EOS detector (Wyatt TechnologyCorp. Santa Barbara, CA). The refractive index was measured using a RI2000 detector (Schambeck SFD). Data analysis was performed with the ASTRA software (Wyatt TechnologyCorp. Santa Barbara, CA). The averaged molecular weight represents the measurements across the elution peak.

X–ray scattering data were collected using an inline HPLC setup, at the Bio–SAXS beamline (BM29) of the European Synchrotron Radiation Facility. Inline size–exclusion chromatography was performed at a temperature of 10°C using a Superdex Increase 200 10/300 GL column equilibrated in SEC buffer. Data were collected with a photon–counting Pilatus 1M detector at a sample–detector distance of 2.86 m, a wavelength of λ = 0.991 Å and an exposure time of 1 second/frame. A momentum transfer range of 0.008 to 0.47 Å^−1^ was covered (q = 4π sinθ/λ, where θ is the scattering angle and λ the X–ray wavelength). Data collected across the peak were subtracted from buffer scattering and the frames 1904 to 2264 showing a constant radius of gyration (Rg) were merged for further analysis. Rg values were obtained from the Guinier approximation sRg < 1.3 using Primus^46^. Distance distribution functions p(r) and the Porod volumes Vp were computed from the entire scattering curve using GNOM^46^. CORAL from the ATSAS suite was used to model the STAT2_133–679_–IRF9–IAD complex using the STAT2_133–_ 679:IRF9–IAD homology model and the IRF9–IAD structure as the input files. The final model conforms well to the scattering data with a chi^2^=1.64. The model for the ISGF3 complex bound to a ISRE DNA sequence 5’-GGGAAATGGAAACT-3’ was obtained by positioning a STAT1 dimer bound to a GAS DNA sequence (PDB; 1BG5) on the first 5’-GGAA-3’ repeat. A homology model of the IRF9 –DBD was positioned on the second GAAA repeat based on the structure of DNA-bound IRF3 (PDB; 1T2K). The final model was obtained by overlaying the STAT2-IRF9 complex onto the distal copy of the STAT1 dimer.

### Crosslinking mass spectrometry

Crosslinking was performed using the STAT2_133–679_/IRF9–IAD complex by incubating with isotope–labelled disuccinimidyl suberate (DSS) as described previously^47^. Protein digestion was performed at 37°C with LysC for 4 hrs followed by trypsin digestion overnight; digested peptides were enriched by SEC. Fractions were injected onto a nanoAcquity ultraperformance liquid chromatography column connected to a LTQ Orbitrap Velos Pro instrument (Thermo Scientific) for liquid chromatography based mass spectrometry measurements. Data processing was performed using xQuest/xProphet. Identified crosslinks were mapped onto the SAXS model of the STAT2_133–679_/IRF9–IAD heterodimer and analysed using Xlink analyzer^48^.

### Isothermal titration calorimetry

Proteins were extensively dialysed against ITC buffer (20 mM Hepes pH 7.5, 300 mM NaCl, 2% glycerol) and subjected to calorimetry using a MicroCal ITC200 system (Malvern Instruments). STAT2_133–679_ at 30 μM, was titrated with different IRF9–IAD variants at concentrations ranging from 150 μM to 320 μM. For IRF9–IADwt, titrations were carried out by injection of 1.5 μl of IRF9–IAD (ITC200) every 180s into the sample cell containing STAT2_133–679_ variants. For mutant IRF9–IAD, 16 successive injections of 2.5 μl were done every 240s. STAT1_136–713_ variants or STAT2_133–679_ V171E were titrated against IRF9-IAD every 120s for 20 injections. Enthalpy change data for titrations were double background corrected via subtraction of protein into buffer measurement. The data were fit using MicroCal Origin 7.0 software (OriginLab).

### Cell culture

All media and supplements were from Gibco, except where indicated. IRF9^−/−^/IRF−3^−/−^ MEFs, kindly provided by Dr. K. Mossman, McMaster University, Hamilton, Canada), were immortalized using the 3T3 protocol and cultured in MEM medium supplemented with non– essential amino acid, sodium pyruvate, 1% L-glutamine and 10% heat-inactivated fetal bovine serum (HI–FBS). HEK293 cells (ATCC) were cultured in DMEM medium containing 1% L-glutamine and 10% Fetalclone III (Hyclone). All cultures were performed without antibiotics and controlled for the absence of mycoplasma contamination using the MycoAlert Mycoplasma Detection Kit (Lonza).

### Luciferase reporter assays

MEF cells were cotransfected with the pRL–null Renilla (*Renilla* luciferase, internal control), the *IFIT1*prom–pGL3 *firefly* luciferase reporter ^49^ and the indicated Flag–IRF9wt or mutant expression plasmid using the *Trans*IT–LT1 transfection reagent (Mirus). At 8h post–transfection, cells were stimulated for 16h with 200U/mL murine IFNβ (PBL Assay Science) Luciferase activities were quantified using the dual luciferase reporter assay kit (Promega). Relative luciferase activities were calculated as the *firefly* luciferase/*Renilla* ratio. Protein extracts were subjected to SDS–PAGE electrophoresis and analysed by immunoblot using the anti–Flag M2 (F1804, Sigma–Aldrich) and anti–actin (A5441, Sigma–Aldrich) antibodies. Immunoreactive bands were visualized using the Western Lightning Chemiluminescence Reagent Plus (Perkin Elmer Life Sciences) acquired on an ImageQuant LAS 4000mini apparatus (GE Healthcare).

### Co-immunoprecipitation

HEK293 cells were transfected with FLAG-IRF9wt or mutants together with HA-STAT2 encoding plasmids using the calcium phosphate method. Cells were lysed by sonication in 50 mM Hepes pH 7.4, 150 mM NaCl, 5 mM EDTA, 10% Glycerol, 1% Triton, 10 μg/mL aprotinin, 10 μg/mL leupeptin, 5 mM NaF, 1 mM activated Na_3_VO_4_, 2 mM p-nitrophenyl phosphate and 10 mM β-glycerophosphate pH 7.5. Cell lysates (1mg) were subjected to immunoprecipitation using 2 μg anti-Flag M2 antibodies (F1804, Sigma-Aldrich) for 3h at 4 °C. Elution of immunocomplexes was performed by incubation on ice for 2h in lysis buffer containing 100 μg/ml FLAG-peptide (F3290, Sigma-Aldrich). Immunocomplexes were analyzed by immunoblot using the anti-FLAG M2 (F1804, Sigma-Aldrich), anti-HA (ab9110, Abcam) and anti-actin (A5441, Sigma-Aldrich) antibodies as described above.

### Data availability

Coordinates for the IRF9–IAD and the IRF9–STAT2 complex are available from the Protein Data Bank under accession number xxx and xxx, respectively.

## Acknowledgements

This work was supported by a grant from the Natural Sciences and Engineering Research Council of Canada (NSERC, #355306) to NG. SR is a fellow of Foundation ARC pour la recherche sur le Cancer, France. NG is recipient of a Research Chair in signaling in virus infection and oncogenesis from Université de Montréal. We thank the staff at the ESRF beamlines BM29, ID30a–1,3 (MASSIF) for their support during data collection. We thank Joanna Kirkpatrick and the proteomic core facility at EMBL for processing and analysis of crosslinked samples. We thank the Partnership for Structural Biology (Grenoble) for providing access to their biophysical platform. We thank Dr. K. Mossman (McMaster University) for the MEF cells used in this study and Dr. Thomas Decker (Max F. Perutz Laboratories, University of Vienna) for comments on the manuscript.

## Author contributions

S.R., S.G, J.D. and E.C. performed the experiments. S.R., E.C., N.G., and D.P. designed experiments and analysed data. S.R. and D.P. wrote the manuscript.

